# Neural signatures of spontaneous transitions between internal and external thought

**DOI:** 10.64898/2026.08.27.747628

**Authors:** Mengqi Zhao, Mengting Zhang, Haowen Su, Philip R. Liu, Hongmi Lee

## Abstract

The human mind constantly shifts between internal representations and the external environment, yet the neural mechanisms underlying such spontaneous transitions remain underexplored. Here, we analyzed a think-aloud functional magnetic resonance imaging dataset, in which participants continuously verbalized their thoughts, to identify neural activity predicting transitions between internally and externally oriented thought. Internal-to-external transitions were preceded by increased activation in the salience/ventral attention network, with the strongest effect observed in the right temporoparietal junction. The spatial pattern of this pre-transition activation was positively associated with acetylcholine receptor density, suggesting a role for cholinergic signaling in cognitive reorientation. Pre-transition activation was itself preceded by a large-scale brain state proposed to serve as a flexible hub between functionally specialized states, indicating that spontaneous transitions are more likely when the brain occupies this intermediate configuration. Together, these findings suggest that multilevel neural mechanisms support flexible reorientation along the internal-external dimension of spontaneous cognition.

## INTRODUCTION

In daily life, we may drift into a daydream in the middle of a conversation, only to have another person’s words suddenly draw us back. Such transitions between engagement with the external environment and internally generated representations are a ubiquitous feature of everyday mental life and are important for adaptive cognition, allowing us to balance information acquisition, memory retrieval, decision making, and future planning^1–3^. Prior research has examined these internal and external modes of cognition from several complementary perspectives. In attention research, they are often distinguished by the target of limited attentional resources: external attention selects and filters physically presented stimuli, whereas internal attention prioritizes representations maintained in working memory^3–5^. Similarly, memory research has distinguished encoding-oriented states that facilitate the processing of external information from retrieval-oriented states that support the reinstatement of internally stored information^6–8^. More broadly, studies of large-scale brain states suggest that cognition naturally alternates between externally and internally oriented modes over timescales ranging from milliseconds to minutes^9–12^.

Despite differences in emphasis and terminology, these perspectives converge on the view that internally and externally oriented cognition can either cooperate or compete depending on the current context^13^, and that they are supported by partially distinct neural mechanisms^1,14^. Internally oriented cognition has been consistently linked to the default mode network (DMN), including medial temporal and posterior parietal regions that support internally generated representations such as memory retrieval^1,9,15^. In contrast, externally oriented cognition relies more heavily on sensory, dorsal attention, and frontoparietal control networks involved in the processing of perceptual information^3,16^. Alongside these large-scale functional networks, neuromodulatory systems provide a complementary neurochemical mechanism for regulating the balance between external and internal processing. The cholinergic system has been proposed as a key candidate^9,14^, as acetylcholine is thought to enhance afferent sensory input relative to internal feedback processing, thereby promoting the encoding of external information ^17,18^. Consistent with this account, nicotinic cholinergic stimulation has been shown to enhance performance on hippocampus-mediated tasks requiring externally oriented attention and perception^19^, and functional connectivity between the cholinergic basal forebrain and hippocampus is stronger during externally guided than memory-guided attention^2^.

These neural mechanisms underlying external and internal modes of cognition have been characterized primarily using task-based designs in which transitions between modes, as well as their timing, are controlled by the experimenter. For example, participants are explicitly cued to engage in externally oriented processes, such as making judgments about perceptual objects or features^2,20,21^, or internally oriented processes, such as maintaining working memory representations or retrieving information from long-term memory^20,21^. A complementary line of mind wandering research has examined more spontaneous transitions in the absence of externally imposed cues, using intermittent thought probes or experience sampling methods, in which participants are periodically prompted to report the focus of their attention^22,23^. A self-caught meta-awareness paradigm has also been used, in which participants report episodes of attentional wandering during ongoing internally or externally focused tasks whenever they become aware of them^24^. However, these intermittent reports cannot capture cognitive states or transitions occurring between reports, nor can they reliably determine the timing of those transitions. Moreover, the reporting process itself may disrupt the ongoing stream of thought and alter the associated neural responses. Consequently, the neural dynamics that predict and potentially trigger naturally occurring transitions between internal and external modes of cognition remain relatively underexplored, despite their ubiquity in everyday life and their central role in theories of large-scale brain states^3,9,14^.

To characterize the neural signatures preceding spontaneous transitions between internally and externally oriented cognition, the present study reanalyzed a publicly available functional magnetic resonance imaging (fMRI) dataset collected using a think-aloud paradigm^25,26^. The think-aloud paradigm, in which participants continuously verbalize their ongoing thoughts, is widely used to study the dynamics of spontaneous thought^27–29^ and provides an ideal approach for examining naturally occurring cognitive and neural state transitions^25,30–32^. Specifically, the explicit verbalization of mental states allows us to determine whether participants are in an internal or external mode without interrupting the continuous flow of thought. It also enables the timing of transitions between the two modes to be identified, allowing neural responses to be aligned to transition onset and examined in the period preceding each transition. Additionally, because think-aloud responses are typically collected in a stable environment without experimentally imposed stimulus changes^27,28,30,31^, this paradigm makes it possible to examine almost purely spontaneous switching between internal and external modes.

Specifically, we addressed three questions. First, we examined which cortical regions and functional brain networks are engaged before spontaneous transitions from internally to externally oriented thought, compared with transitions between thoughts within the same mode. We hypothesized that internal-to-external transitions would be preceded by greater activation in cognitive reorienting systems, particularly the salience/ventral attention network, which has been implicated in detecting salient external information and reallocating attentional resources accordingly^33,34^. Second, we tested whether the neural activity preceding internal-to-external transitions is related to the spatial organization of neurotransmitter systems. To address this question, we compared the whole-brain spatial distribution of pre-transition activation with maps of neurotransmitter receptor density derived from existing positron emission tomography (PET) datasets^35–42^. Based on prior evidence implicating acetylcholine in externally oriented processing^2,17,19^, we hypothesized that cholinergic receptor density would be preferentially associated with pre-transition activation during internal-to-external transitions. Finally, we investigated whether spontaneous transitions between internal and external thought are preceded by large-scale brain-state reconfigurations, building on evidence that spontaneous neural activity is organized into recurring large-scale brain states that vary in attentional engagement and cognitive demand^31,43,44^. To test this possibility, we correlated whole-brain activation patterns preceding transitions with brain-state templates previously derived from task and resting-state fMRI data^44^. We hypothesized that spontaneous internal-to-external transitions would be preceded by increased similarity to an intermediate, or "base," brain state that bridges functionally specialized states^44,45^, supporting flexible switching between internally and externally oriented modes of cognition.

## RESULTS

### Spontaneous transitions between internal and external thoughts occur during think-aloud

To characterize the neural signatures preceding spontaneous transitions between internally and externally oriented thoughts, we reanalyzed a think-aloud fMRI dataset originally collected for our previous study^25^. Participants continuously verbalized their thoughts for 10 minutes while external sensory stimuli (a fixation cross and scanner noise) remained constant. The speech recordings were transcribed, and each sentence was classified as either an internally oriented or an externally oriented thought (Fig. 1a). Across the 75 participants with usable fMRI data, participants produced an average of 102.32 sentences (SD = 35.52), excluding filler utterances. Although participants primarily produced internal thoughts (M = 93.08%, SD = 9.26), external thoughts also occurred during the task (M = 6.92%, SD = 9.26), allowing us to identify all four types of spontaneous transitions between thought orientations: internal-to-internal (M = 89.37%, SD = 13.43), internal-to-external (M = 3.72%, SD = 4.33), external-to-internal (M = 4.05%, SD = 4.54), and external-to-external (M = 2.86%, SD = 5.36).

**Fig. 1.**
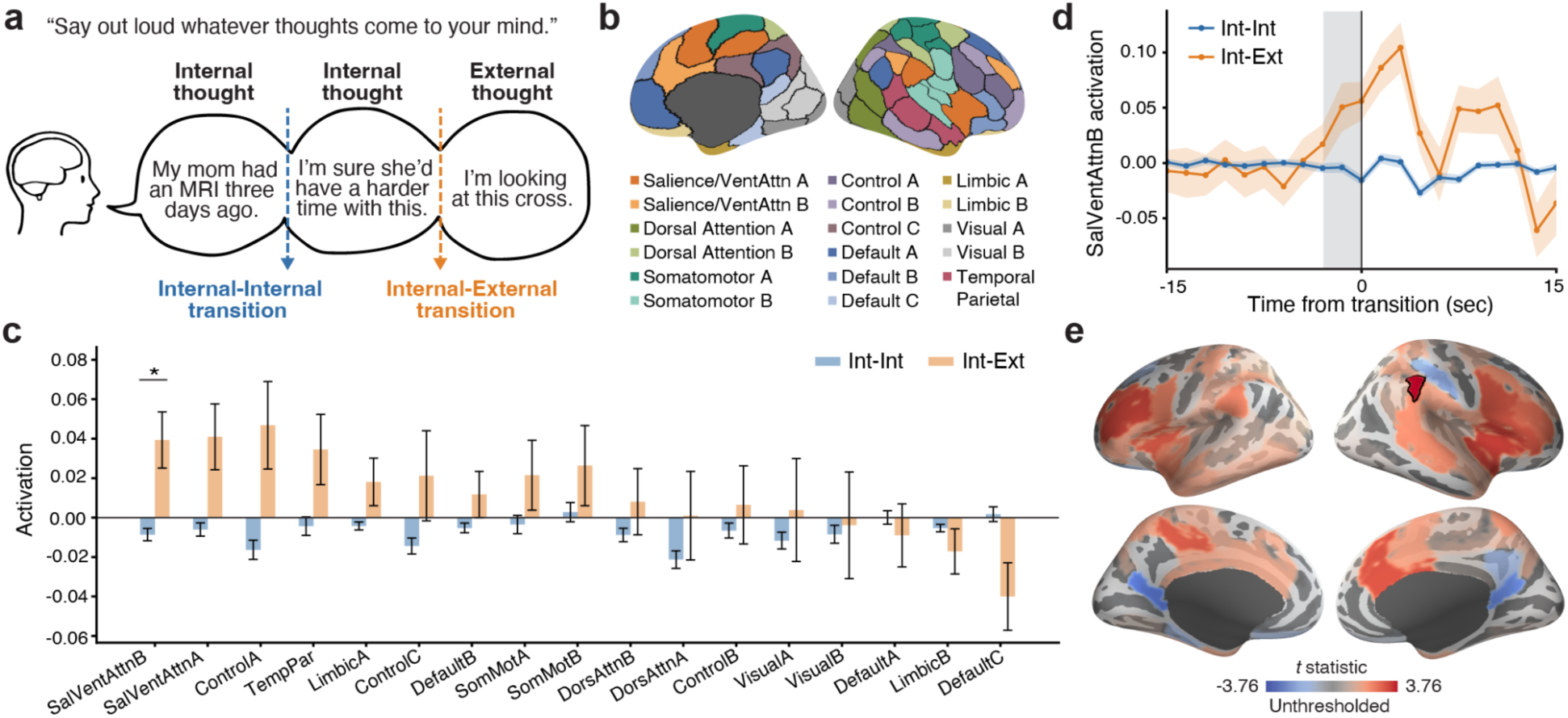
Neural activation preceding internal-to-external thought transition. (**a**) Schematic of the 10-min think-aloud task and identification of thought transitions. Participants continuously verbalized their ongoing thoughts while undergoing fMRI. Speech recordings were transcribed and segmented into sentences. Each sentence was classified as internally or externally oriented, allowing consecutive sentence boundaries to be categorized as internal-to-internal or internal-to-external thought transitions (external-to-internal and external-to-external boundaries were also possible but are not shown in the schematic). (**b**) Cortical parcellation based on the Schaefer 100-parcel, 17-network atlas^46^, shown on the medial (left) and lateral (right) surfaces of the right hemisphere. (**c**) Mean activation during the pre-transition analysis window (-2 to 0 TRs relative to the transition) across the 17 networks. Blue (internal-to-internal; Int-Int) and orange (internal-to-external; Int-Ext) bars represent the group mean for each transition type (N = 54), and error bars indicate the SEM. The asterisk indicates a significant difference between transition types, as determined by two-tailed paired *t*-tests with FDR correction (*q* < 0.05). (**d**) Mean activation time courses in salience/ventral attention network B aligned at thought transition. Time zero denotes the onset of post-transition thoughts, and the gray shaded region indicates the pre-transition analysis window. Solid lines and circles indicate the mean activation across participants (N = 56) at each time point, and shaded areas indicate the SEM. (**e**) Whole-brain maps for the pre-transition internal-to-external versus internal-to-internal contrast. Red regions indicate greater activation preceding internal-to-external transitions. Maps show unthresholded *t* statistics, and the parcel outlined in black indicates the right temporoparietal junction parcel that survived FDR correction (*q* < 0.05). SalVentAtten, Salience/ventral attention; TempPar, Temporal parietal; SomMot, Somatomotor; DorsAttn, Dorsal attention.

Because the majority of reported thoughts were internally oriented, the duration of consecutive episodes (i.e., dwell times) of externally oriented thoughts were relatively brief (M = 8.56 s, SD = 6.46) compared with those of internally oriented thoughts (M = 263.18 s, SD = 210.67). As a result, external-to-internal transitions typically occurred shortly after the preceding internal-to-external transitions, making it difficult to dissociate neural activity preceding external-to-internal transitions from carryover hemodynamic responses elicited by the preceding internal-to-external transitions (Supplementary Fig. 1). We therefore focused all subsequent analyses on comparing internal-to-external with internal-to-internal transitions. Participants who did not exhibit any internal-to-external transitions were excluded, leaving 56 participants for analysis.

After this exclusion, 4.98% of transitions were internal-to-external (SD = 4.34), and 85.84 % were internal-to-internal (SD = 13.88).

### Salience/ventral attention network activation precedes internal-to-external thought transition

We first examined which functional brain networks are engaged before spontaneous transitions from internally oriented to externally oriented thought. For each of the 17 functional networks defined by the Schaefer atlas^46^ (Fig. 1b), we computed the mean activation during a 4.5-s pre-transition window comprising the two TRs immediately preceding each transition and the transition onset TR, separately for internal-to-external and internal-to-internal transitions. The window was selected based on prior studies reporting neural changes approximately 1-5 s before transitions in thought and awareness^31,47,48^. Paired-samples *t*-tests comparing the two transition types revealed significantly greater pre-transition activation in salience/ventral attention network B before internal-to-external than internal-to-internal transitions after correction for multiple comparisons (*t*(53) = 3.27, *p* = 0.002, 95% CI = [0.02, 0.08], Cohen’s *d* = 0.44, Figs. 1c-d). Salience/ventral attention network A, control network A, and the temporal parietal network showed similar but weaker effects that did not survive multiple-comparison correction. Supplementary Table 1 provides detailed statistics for all functional networks.

To more precisely localize the effects of transition type on pre-transition activation, we next conducted a parcel-level analysis across the 100 cortical parcels spanning the entire cerebral cortex, as defined by the Schaefer atlas (Fig. 1e). Only one parcel survived correction for multiple comparisons: a parcel in the right inferior parietal lobule overlapping the right temporoparietal junction (TPJ), which showed greater pre-transition activation before internal-to-external than internal-to-internal transitions (*t*(53) = 3.76, *p* < 0.001, 95% CI = [0.04, 0.15], Cohen’s *d* = 0.51). Detailed parcel-level statistics are provided in Supplementary Table 2.

Together, these findings suggest that the salience/ventral attention network, particularly the right TPJ, contributes to spontaneous transitions from internally to externally oriented thought.

### Pre-transition salience/ventral attention network activation cannot be explained primarily by semantic shifts

Previous studies have shown that semantic relatedness is a major organizing principle of spontaneous thought^25,29,49^, and that larger semantic shifts between consecutive thoughts elicit stronger neural responses following thought boundaries^25^. We therefore tested whether the increased salience/ventral attention network activation preceding internal-to-external transitions could be explained by larger semantic shifts between internal and external thoughts than between successive internal thoughts.

To quantify semantic similarity between pre- and post-transition thoughts, each sentence was converted into a 768-dimensional embedding vector using a pretrained Sentence Transformers language model^50^. Semantic similarity was then defined as the cosine similarity between the embedding vectors of consecutive sentences, following the procedures used in our previous study^25^ (Fig. 2a). We found that internal-to-external transitions indeed showed lower semantic similarity between consecutive thoughts than internal-to-internal transitions (*t*(53) = -8.11, *p* < 0.001, 95% CI = [-0.14, -0.09], Cohen’s *d* = -1.1, Fig. 2b). To control for this difference, we selected, within each participant, a subset of internal-to-internal transitions with the lowest semantic similarity between pre- and post-transition thoughts until their mean semantic similarity matched that of the internal-to-external transitions. Following this matching procedure, semantic similarity no longer differed between the two transition types (*t*(53) = 1.5, *p* = 0.14, 95% CI = [-0.003, 0.02], Cohen’s *d* = 0.2, Fig. 2b).

**Fig. 2.**
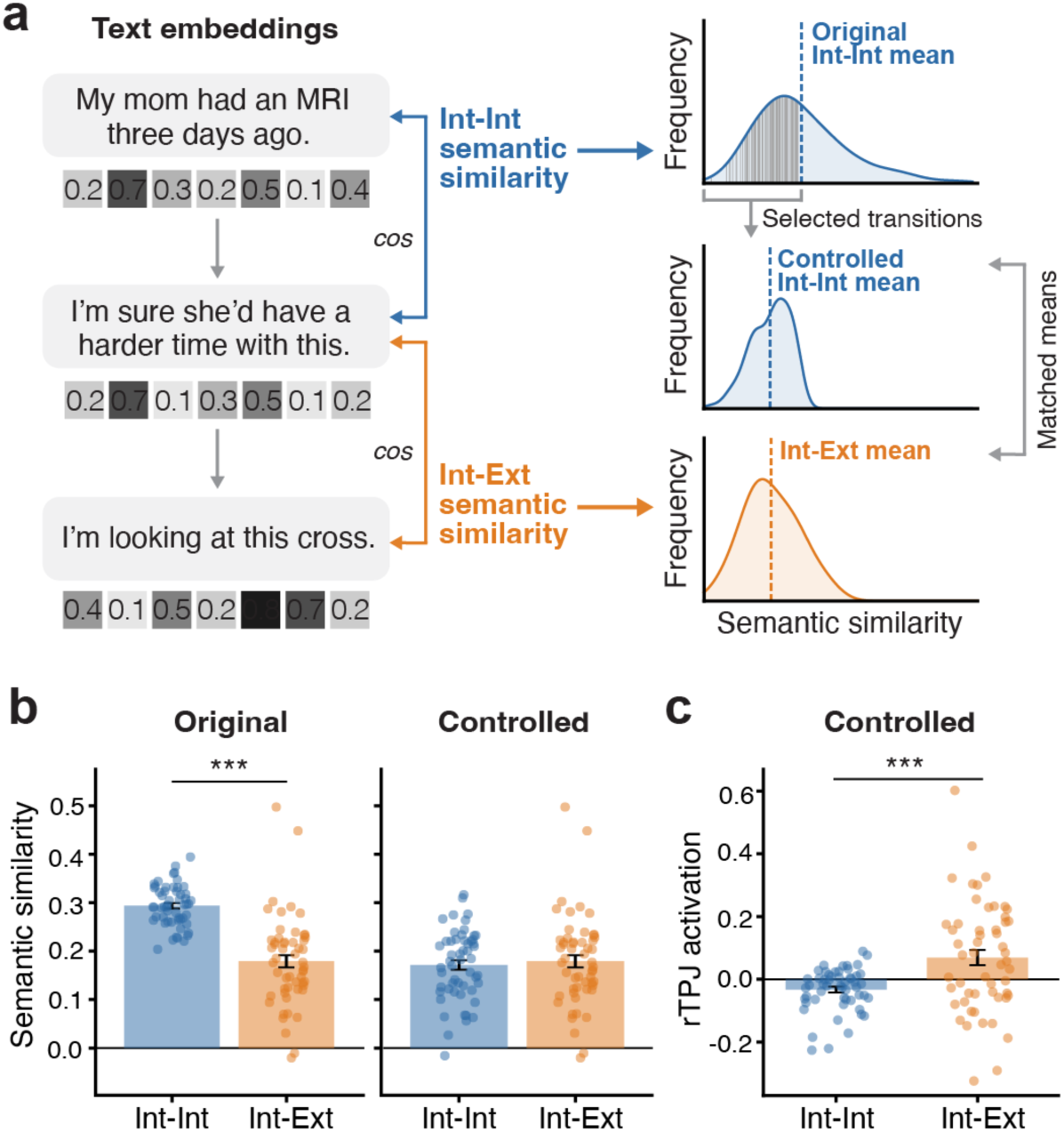
Controlling for semantic similarity at thought transitions. (**a**) Schematic of the semantic-similarity control analysis. Each thought unit was converted into a text-embedding vector, and semantic similarity between consecutive thoughts was quantified using cosine similarity. A subset of internal-to-internal transitions with the lowest semantic similarity was selected to match the mean semantic similarity of the internal-to-external transitions. (**b**) Semantic similarity before matching (Original; left panel) and after matching (Controlled; right panel). (**c**) Pre-transition activation in the right temporoparietal junction parcel (rTPJ) after controlling for semantic similarity. In **b** and **c**, each dot represents one participant (N = 54); bars indicate the mean across participants, and error bars show the SEM. \*\*\**p* < 0.001. Int-Int, internal-to-internal; Int-Ext, internal-to-external.

We next repeated the pre-transition activation analysis using the semantically matched subset of internal-to-internal transitions. The results remained largely unchanged: salience/ventral attention network B continued to show greater pre-transition activation for internal-to-external than internal-to-internal transitions (*t*(53) = 3.22, *p* = 0.002, 95% CI = [0.02, 0.08], Cohen’s *d* = 0.44), and the right TPJ parcel likewise showed a significant effect of transition type (*t*(53) = 3.75, *p* < 0.001, 95% CI = [0.05, 0.16], Cohen’s *d* = 0.51, Fig. 2c).

Detailed statistics for all functional networks and suprathreshold cortical parcels following semantic matching are reported in Supplementary Tables 3 and 4. These findings suggest that differences in semantic similarity alone are unlikely to account for the increased pre-transition activation preceding internal-to-external transitions. Instead, the effect appears to reflect the shift from an internally to an externally oriented mode of cognition.

### Eye movements increase before transitions to visually oriented external thoughts

In addition to changes in verbal expression, spontaneous thought shifts and attentional reorienting may also be accompanied by changes in eye movements^51,52^, which could in turn influence neural activity^53^. To examine this possibility, we used DeepMReye^54^, an MRI-based eye-tracking method, to estimate gaze position directly from the fMRI data. We then computed TR-by-TR gaze displacement (vector length) as an approximate measure of eye movement amplitude (Fig. 3a). Consistent with previous studies using the same approach^55,56^, a linear mixed-effects model showed that the mean gaze vector length during each sentence predicted visual cortical activation (Supplementary Fig. 2), supporting the validity of the MRI-derived gaze vector length measure.

**Fig. 3.**
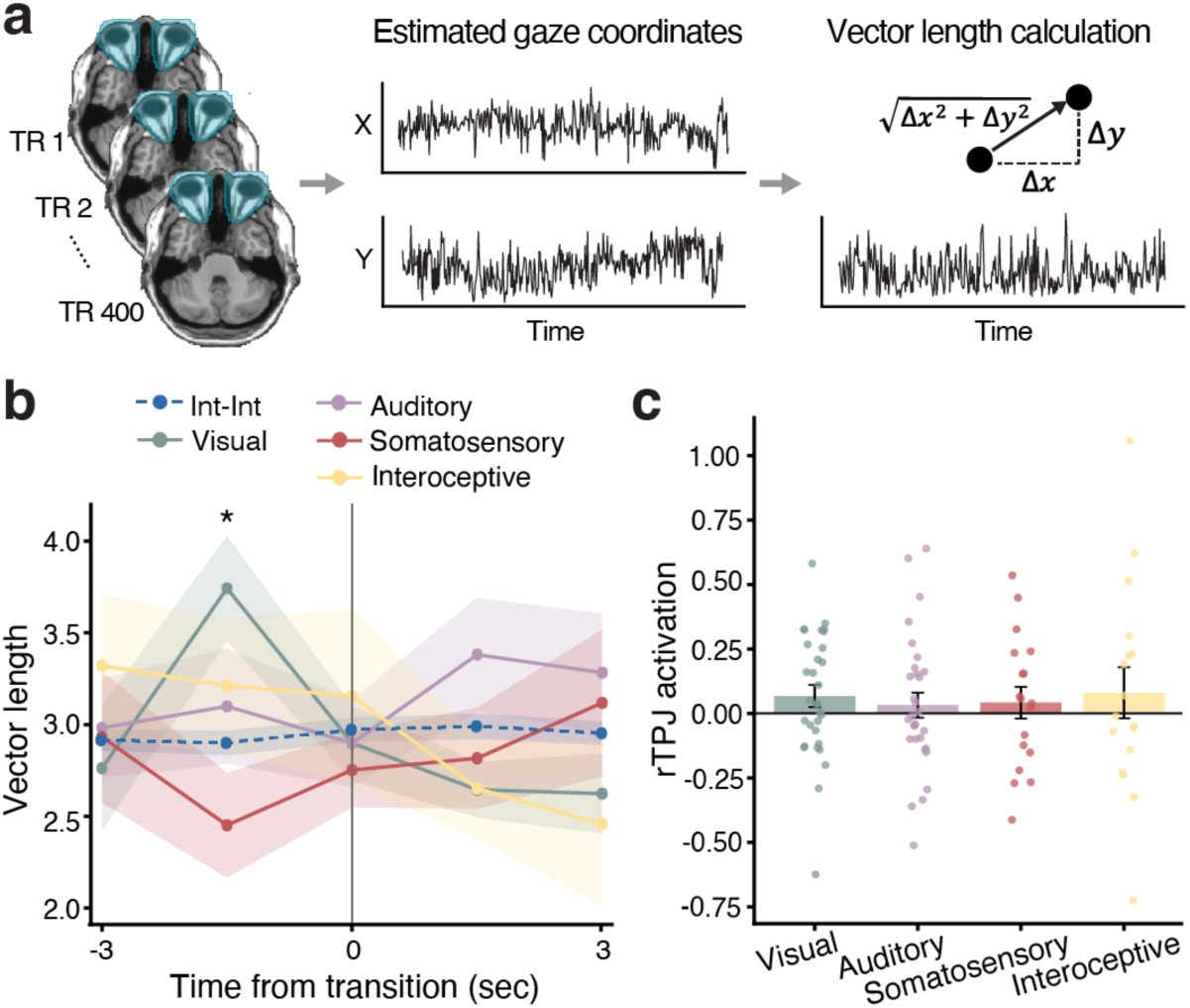
Pre-transition eye movements and neural activation across sensory modalities. (**a**) Schematic of the fMRI-derived eye-movement analysis. DeepMReye^54^ was used to estimate the horizontal and vertical gaze position at each TR, and eye-movement vector length was calculated as the Euclidean distance between consecutive gaze positions. (**b**) Mean vector length time courses aligned at different types of thought transitions (blue = internal-to-internal, green = internal-to-visual, purple = internal-to-auditory, red = internal-to-somatosensory, and yellow = internal-to-interoceptive). Time zero denotes the onset of post-transition thoughts. Colored lines and circles indicate the mean vector length at each time point across participants (N = 56, 31, 29, 18, and 17 for internal-to-internal, visual, auditory, somatosensory, and interoceptive internal-to-external transitions, respectively). Shaded areas indicate the SEM. The asterisk at 1.5 s before the transition indicates a significant difference between the internal-to-internal and internal-to-visual conditions after FDR correction (*q* < 0.05). (**c**) Mean pre-transition activation in the right temporoparietal junction (rTPJ) parcel for internal-to-external transitions, separated by sensory modality. Individual dots represent participants (N = 31, 29, 18, and 17 for vision, audition, somatosensation, and interoception, respectively). Bars indicate the group mean, and error bars show the SEM. Int-Int, internal-to-internal.

To test whether eye movements differed between internal-to-external and internal-to-internal transitions, we examined gaze vector length within a 5-TR (7.5-s) window centered on the transition onset. A repeated-measures ANOVA with transition type and time points as within-subject factors revealed a significant interaction between the two factors (*F*(4, 204) = 2.84, *p* = 0.025, *η*^2^ = 0.01), with no significant main effects of transition type (*F*(1, 51) = 0.33, *p* = 0.57, *η*^2^

= 0.001) or time point (*F*(4, 204) = 1.49, *p* = 0.206, *η*^2^ = 0.01). Post hoc analyses showed that gaze vector length was significantly greater 1 TR before internal-to-external than before internal-to-internal transitions, which remained significant after correction for multiple comparisons across time points (internal-to-external *M* = 3.34, internal-to-internal *M* = 2.9, *t*(52) = 2.76, *p* = 0.008, 95% CI = [0.12, 0.76], Cohen’s *d* = 0.38).

We next tested whether the increase in gaze vector length 1 TR before transition onset depended on the sensory modality of the post-transition external thought, specifically whether it was driven by transitions to visual content (Fig. 3b). Internal-to-external transitions were classified as visual, auditory, somatosensory, or interoceptive based on the sensory modality expressed in the post-transition thought. Paired comparisons with internal-to-internal transitions showed that only visually oriented internal-to-external transitions were associated with significantly greater pre-transition gaze vector length after correction for multiple comparisons across sensory modalities (*t*(30) = 3.33, *p* = 0.002, 95% CI = [0.36, 1.5], Cohen’s *d* = 0.6). No significant effects were observed for auditory (*t*(28) = 0.51, *p* = 0.612, 95% CI = [-0.41, 0.68], Cohen’s *d* = 0.1), somatosensory (*t*(17) = -1.58, *p* = 0.132, 95% CI = [-1.08, 0.15], Cohen’s *d* = -0.37), or interoceptive transitions (*t*(16) = 0.32, *p* = 0.753, 95% CI = [-0.71, 0.97], Cohen’s *d* = 0.08).

### Pre-transition salience/ventral attention network activation is general across sensory modalities

Our MRI-based eye-movement analysis suggested that the pre-transition oculomotor changes may be specifically related to visual processing. Moreover, because processing of different sensory modalities engages partially distinct brain regions^57^, it is possible that the observed pre-transition activation reflects modality-specific sensory preparation rather than a general shift toward externally oriented thought. To test this possibility, we categorized post-transition external thoughts as visual, auditory, somatosensory, or interoceptive using the same procedures as in the eye movement analysis. We then examined whether pre-transition activation during internal-to-external transitions varied as a function of the sensory modality of the upcoming external thought (Fig. 3c). However, pre-transition activation in salience/ventral attention network B did not differ significantly between any pair of sensory modalities (all *p*s > .14). Likewise, the right TPJ parcel showed no significant differences in pre-transition activation across sensory modalities (all *p*s > .13). Detailed modality-specific statistics are reported in Supplementary Tables 5 and 6.

### Acetylcholine receptor density is positively associated with internal-to-external pre-transition activation

Thus far, we have shown that transitions from internally to externally oriented thoughts are preceded by increased activation in a specific subset of brain regions, particularly within the salience/ventral attention network. This finding raises the question of why these regions are preferentially engaged during the transition. One possibility is that their involvement is related to underlying neurochemical organization. In particular, previous studies have implicated acetylcholine in attentional orienting and the regulation of internally versus externally directed cognition^9,14^. We therefore tested whether brain regions with higher cholinergic receptor density exhibited stronger pre-transition activation during internal-to-external thought transitions (Fig. 4a).

**Fig. 4.**
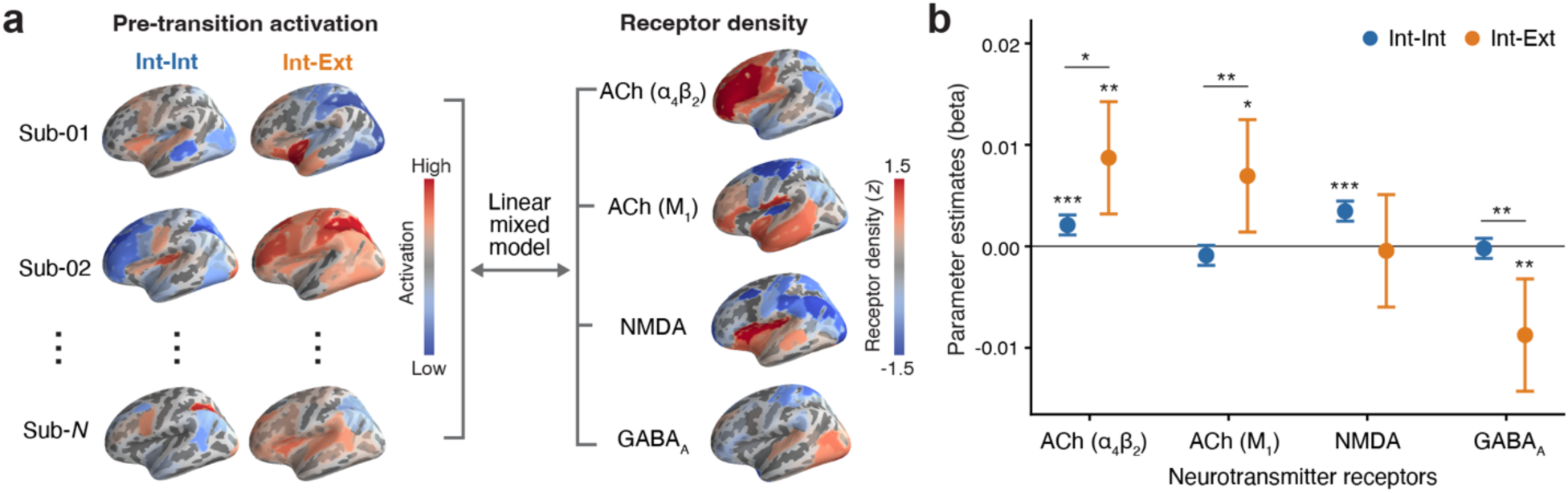
Relationship between neurotransmitter receptor density and pre-transition activation. (**a**) Schematic of the receptor-density analysis. For each participant and transition type, a whole-brain parcel-wise activation map was generated by averaging pre-transition activation across transition events within each of the 100 cortical parcels. Linear mixed-effects models were then used to predict these whole-brain pre-transition activation patterns from parcel-wise neurotransmitter receptor density maps (z-scored across parcels) derived from a publicly available dataset^42^. (**b**) Parameter estimates from linear mixed-effects models predicting pre-transition activation during internal-to-internal and internal-to-external transitions using neurotransmitter receptor density. Circles indicate parameter estimates, and error bars indicate 95% confidence intervals. Statistical significance of receptor-activation associations for each transition type was determined from models fit separately to internal-to-internal and internal-to-external transitions. Differences between transition types were assessed using models that included the interaction between receptor density and transition type. Two-tailed Wald tests were used to test individual fixed effects. \**p* < 0.05, \*\**p* < 0.01, \*\*\**p* < 0.001. Int-Int, internal-to-internal; Int-Ext, internal-to-external; ACh, Acetylcholine.

We focused on two cholinergic receptors: the alpha-4-beta-2 nicotinic acetylcholine receptor (α4β2) and the muscarinic M1 receptor (M1). As comparison receptors, we included the N-methyl-D-aspartate receptor (NMDA) and the gamma-aminobutyric acid type A receptor (GABAA), which mediate major excitatory and inhibitory neurotransmission in the brain, respectively^58,59^. PET-derived receptor density maps for each receptor were obtained from a previously published dataset ^42^. We then used linear mixed-effects models to examine the relationship between parcel-wise receptor density and pre-transition activation.

As expected, when internal-to-internal and internal-to-external transitions were modeled separately (Fig. 4b), higher α4β2 (*β* = 0.007, 95% CI = [0.003, 0.011], *p* = 0.002) and M1 (*β* = 0.006, 95% CI = [0.001, 0.01], *p* = 0.014) receptor densities were each associated with greater pre-transition activation during internal-to-external transitions. In contrast, NMDA receptor density was not significantly associated with internal-to-external pre-transition activation (*β* = -0.0004, 95% CI = [-0.005, 0.004], *p* = 0.874), whereas GABAA receptor density showed a negative association (*β* = -0.007, 95% CI = [-0.011, -0.003], *p* = 0.002). For internal-to-internal transitions, higher α4β2 (*β* = 0.002, 95% CI = [0.001, 0.002], *p* < 0.001) and NMDA (*β* = 0.003, 95% CI = [0.002, 0.004], *p* < 0.001) receptor densities were associated with greater pre-transition activation, whereas M1 (*β* = -0.001, 95% CI = [-0.002, 0.0001], *p* = 0.076) and GABAA (*β* = -0.0002, 95% CI = [-0.001, 0.001], *p* = 0.069) receptor densities were not significantly associated with pre-transition activation.

Models including receptor density, transition type, and their interaction showed that the positive associations between receptor density and pre-transition activation were significantly stronger for internal-to-external than for internal-to-internal transitions for both α4β2 (*β* = 0.005, 95% CI = [0.001, 0.01], *p* = 0.026) and M1 (*β* = 0.006, 95% CI = [0.002, 0.011], *p* = 0.008) receptors. In contrast, GABAA receptor density showed a significantly stronger negative association with pre-transition activation for internal-to-external compared with internal-to-internal transitions (*β* = -0.007, 95% CI = [-0.012, -0.002], *p* = 0.004). Furthermore, in models including multiple receptors as simultaneous predictors, α4β2 (*β* = 0.006, 95% CI = [0.001, 0.011], *p* = 0.018) and M1 (*β* = 0.006, 95% CI = [0.001, 0.01], *p* = 0.01) receptor densities remained significant positive predictors of internal-to-external pre-transition activation after controlling for NMDA and GABAA receptor densities. Detailed statistics for the interaction and multiple receptor models are provided in Supplementary Tables 7 and 8. Together, these findings implicate the cholinergic system in the neural processes preceding spontaneous transitions from internal to external modes of cognition.

### Large-scale brain state changes are associated with internal-to-external thought transition

The pre-transition activation in the salience/ventral attention network during internal-to-external transitions suggests neural activity associated with attentional reorienting toward external information. However, the external sensory environment remained essentially unchanged throughout the think-aloud task, particularly with respect to the visual (the scanner bore and gray screen) and auditory (scanner noise) stimuli that most frequently constituted external thoughts. This raises the question of what caused attention to be occasionally redirected to the external environment at these specific moments of transition.

One possibility is that internal-to-external transitions are preceded by a reconfiguration of large-scale brain states. Previous studies have shown that the brain spontaneously alternates between states associated with internally and externally oriented processing, with intermediate states linking the two^12,44,45^. We therefore hypothesized that, before attention is redirected outward, the brain enters an intermediate or transition-like state that facilitates switching between the two modes of cognition. Alternatively, the brain has already shifted toward an externally oriented state, allowing ongoing processing of external sensory input to gradually accumulate until it captures attention.

To test these hypotheses, we compared whole-brain activation patterns preceding internal-to-internal and internal-to-external transitions with large-scale brain-state templates identified in a previous study^44^. That study characterized four recurring brain states: a default mode network (DMN)-dominant state associated with internally oriented processing, a dorsal attention network (DAN)-dominant state associated with externally oriented attentional engagement, a somatomotor (SM)-dominant state associated with sensorimotor processing, and a base state interpreted as a flexible intermediate state (Fig. 5a). For each participant, we computed the correlation between the whole-brain parcel-wise activation pattern at each TR and each brain-state template across the 10 TRs preceding the transition and the transition-onset TR. We then used repeated-measures ANOVAs with transition type and time point as within-participant factors to test whether similarity to each brain state changed before internal-to-external transitions relative to internal-to-internal transitions, particularly at time points preceding the observed pre-transition activation.

**Fig. 5.**
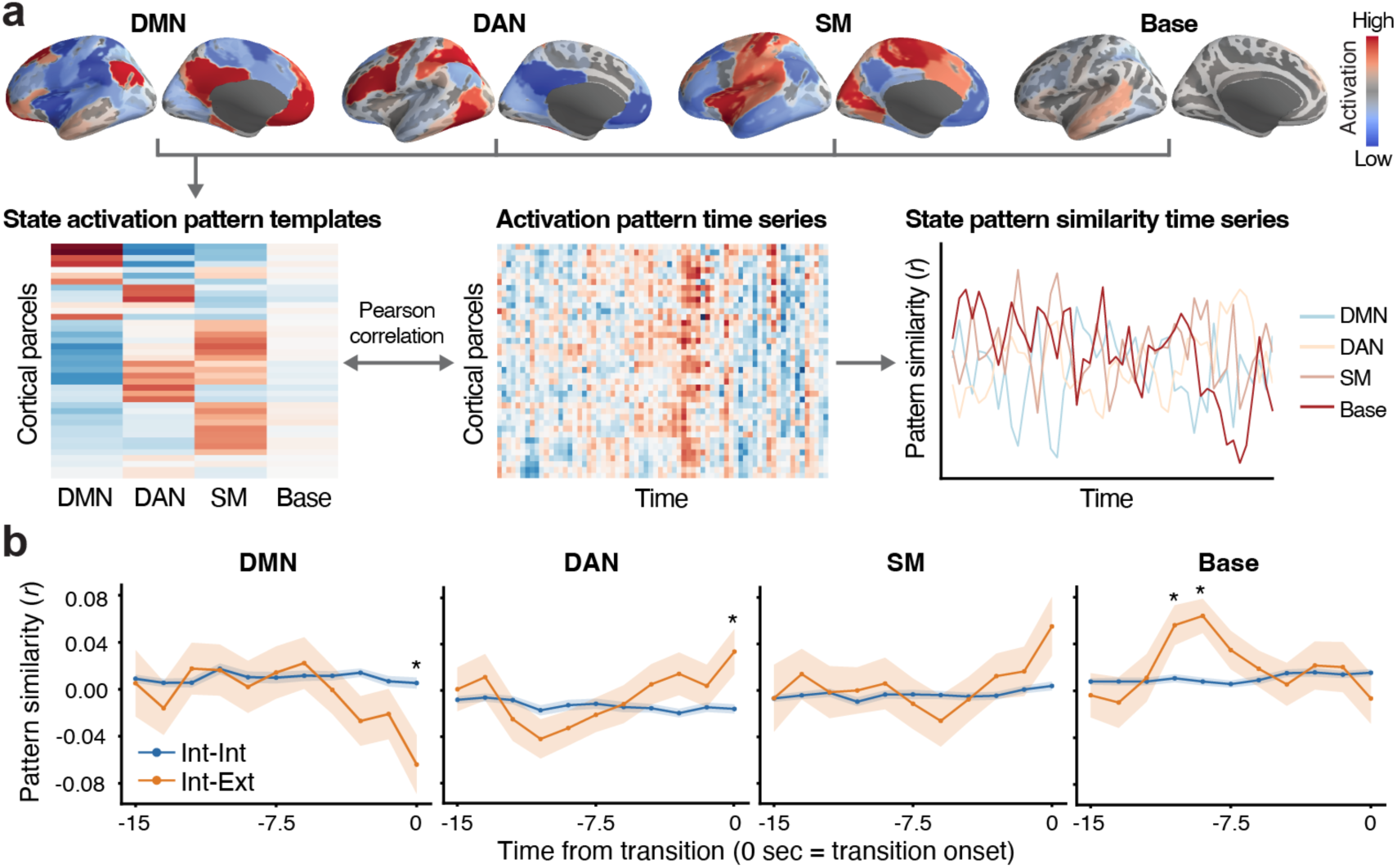
Large-scale brain-state dynamics preceding internal-to-external thought transitions. (**a**) Schematic of the brain-state pattern-similarity analysis. Four whole-brain parcel-wise brain-state template activation patterns were derived from a previous study^44^, corresponding to the default mode network (DMN), dorsal attention network (DAN), somatomotor network (SM), and base states. For each participant, TR-by-TR whole-brain activation patterns were correlated with each brain-state template to generate pattern similarity time courses. Similarity values for the time window preceding thought transitions were then extracted and compared across transition types. (**b**) Mean state-template similarity time courses preceding internal-to-internal (Int-Int) and internal-to-external (Int-Ext) transitions. Time zero denotes the onset of post-transition thoughts. Solid lines and circles indicate the group mean at each time point (N = 56). Shaded areas indicate the SEM. Asterisks mark significant differences between transition types, as determined by two-tailed paired *t*-tests, before correction for multiple comparisons. \**p* < 0.05 (uncorrected).

We found that the base state showed a significant interaction between transition type and time point (*F*(10, 480) = 2.19, *p* = 0.017, *η*^2^ = 0.04), with greater similarity preceding internal-to-external than internal-to-internal transitions at 9 s (*t*(52) = 3.75, *p* < 0.001, 95% CI = [0.03, 0.09], Cohen’s *d* = 0.52) and 10.5 s (*t*(51) = 2.61, *p* = 0.012, 95% CI = [0.01, 0.08], Cohen’s *d* = 0.36) before the transition (Fig. 5b). The DAN state also showed a significant interaction (*F*(10, 480) = 1.94, *p* = 0.038, *η*^2^ = 0.04), although increased similarity emerged later, at the time of the internal-to-external transition (*t*(53) = 2.44, *p* = 0.018, 95% CI = [0.01, 0.09], Cohen’s *d* = 0.33). The DMN state did not show a significant interaction (*F*(10, 480) = 0.93, *p* = 0.508, *η*^2^ = 0.02); however, its similarity decreased at the time of the internal-to-external transition (*t*(53) = -2.66, *p* = 0.01, 95% CI = [-0.12, -0.02], Cohen’s *d* = -0.36). The SM state showed neither a significant interaction (*F*(10, 480) = 0.61, *p* = 0.806, *η*^2^ = 0.01) nor differences between transition types at any time point. After correcting for multiple comparisons across time points, however, only the greater similarity to the base-state at 9 s before the transition remained significant. These results are generally consistent with the hypothesis that the brain enters a transition-like state before the emergence of the salience/ventral attention network activation associated with attentional reorienting.

## DISCUSSION

The present study investigated the neural signatures preceding spontaneous transitions from internally to externally oriented thought. Using a think-aloud fMRI dataset^25,26^, we found that internal-to-external transitions were preceded by increased activation in the salience/ventral attention network, particularly the right TPJ. This pre-transition activation was observed across transitions to different sensory modalities and could not be explained by differences in semantic shifts between consecutive thoughts. Furthermore, the whole-brain spatial pattern of pre-transition activation was positively associated with regional cholinergic receptor density. Finally, these transitions appeared to be preceded by shifts in large-scale brain-state configuration, characterized by increased similarity to the previously identified ‘base state’, a transition hub between more specialized neural states^44^, before the emergence of salience/ventral attention network activation. Together, these findings suggest that spontaneous shifts from internal thought to external engagement are more likely when the brain enters a flexible, transition-prone configuration. This reorientation toward external cognition is then marked by activation in the salience/ventral attention network, potentially modulated by cholinergic signaling.

Because the think-aloud paradigm captured spontaneous, self-initiated thought transitions, we were able to examine the neural activity preceding naturally occurring transitions, rather than responses to externally cued transitions as in previous work^2,3,20^. Compared with internal-to-internal transitions, internal-to-external transitions were associated with greater pre-transition activation in the salience/ventral attention network B, particularly in the right TPJ area. The salience/ventral attention network is well established as a system for detecting salient or unexpected events and reorienting attention^33,34,60^. The right TPJ, a key node of the ventral attention system, has likewise been implicated in stimulus-driven attentional reorienting^60–62^.

Thus, the pre-transition activation observed for internal-to-external transitions may reflect a cognitive reorientation process, potentially involving the reallocation of attentional resources toward external information, that begins before the transition is verbally reported. It is also worth noting that, because semantic similarity between pre- and post-transition thoughts was lower for internal-to-external than internal-to-internal transitions (Fig. 2b), the observed neural activation could alternatively reflect greater semantic or contextual discontinuity between thoughts, as suggested by recent work on event and thought boundaries^25,63^. However, the effect of transition type on pre-transition activation remained significant after controlling for semantic similarity, suggesting that the activation cannot be explained solely by changes in specific thought content. Instead, it may reflect a domain-general marker of cognitive reorientation from internal to external processing.

In parallel with the neural findings, eye-movement amplitude increased shortly before internal-to-external transitions, particularly for transitions toward visually oriented external thoughts. This pattern is unlikely to fully account for the pre-transition neural activation, as activation levels did not differ as a function of the sensory modality of the post-transition thought. Nevertheless, prior work suggests that shifts between internal and external modes of cognition can be accompanied by changes in eye movements, with externally directed cognition often characterized by greater visual scanning and more microsaccades relative to internally directed cognition^64,65^. A higher saccade rate during internal cognition also predicts greater subsequent susceptibility to visual distraction^51^. Therefore, the increased pre-transition eye movement may reflect an early shift away from internal mentation toward sampling the external environment, or a state in which external information becomes more competitive for attention during spontaneous fluctuations between internal and external modes of cognition^66,9^. This interpretation should, however, be treated with caution, as the eye-movement estimates were derived from MRI data^54,55^ and are thus constrained by the method’s limited temporal resolution. Future studies could use conventional high-resolution eye tracking to more precisely characterize oculomotor dynamics around thought transitions, such as microsaccades at sub-second timescales^67^, or pupillary responses indexing arousal and attentional intensity^68,69^.

Another key finding of the present study is that the spatial pattern of pre-transition activation was positively associated with acetylcholine receptor density, especially during internal-to-external transitions. This finding is consistent with a recent theoretical framework proposing that acetylcholine is specifically linked to the internal-external orientation of cognition^14^. Acetylcholine has long been proposed to bias neural processing toward externally oriented cognition by enhancing feedforward sensory input and suppressing internal feedback, thereby promoting the encoding of new information^17,70,71^. Although prior studies of cholinergic effects on memory and cognitive mode switching have focused largely on the hippocampus^2,19,71,72^, our results support the view that acetylcholine’s contribution may extend more broadly across cortical systems^18^. This is also consistent with anatomical evidence that basal forebrain cholinergic neurons project widely throughout the cortex^73,74^, as well as with functional evidence showing cholinergic modulation of cortical activation^75^. Future studies could further integrate measures of cortical and subcortical cholinergic activity to provide a more comprehensive understanding of how the cholinergic system contributes to transitions between cognitive states. Finally, although the present study focused on the cholinergic system, other neuromodulatory systems are also likely to contribute to spontaneous cognitive mode switching. The noradrenergic system, for instance, is a potential candidate given its roles in arousal^76,77^ and large-scale brain state reconfiguration^78,79^. Future work could therefore investigate how multiple neuromodulatory systems interact to support transitions between cognitive states.

Our large-scale brain-state analysis characterized the temporal evolution of whole-brain neural configurations preceding spontaneous thought transitions, motivated by recent empirical and theoretical work suggesting that spontaneous cognition unfolds through recurring and dynamic brain states^12,31^. The observed pattern suggested that internal-to-external pre-transition activation is preceded by an increase in the probability of a base or transition-like state identified in a prior study^44^, followed by an increasing tendency toward an externally oriented dorsal attention (DAN) state and reduced similarity to an internally oriented default mode (DMN) state closer to transition onset. This base state has been characterized in previous research by relatively low global cofluctuation, high global desynchrony, and activity amplitudes close to the global mean^43–45^, and has thus been interpreted as an intermediate or ground state that functions as a flexible transitional hub through which the brain passes when moving between more differentiated network configurations, including the DAN and DMN states^43–45^. Increased similarity to this base state may therefore keep both internally generated representations and external sensory information relatively accessible, increasing the probability of shifting between internally and externally oriented cognition.

If the brain is already in a state that may facilitate switching before transitioning between internally and externally oriented thought, what drives the brain into this transitional state in the first place? One possibility comes from frameworks that conceptualize cognitive transitions as naturally emerging from dynamic competition or relative imbalance between internally and externally oriented processing^9,66^. Under this view, neural systems may cycle across multiple timescales between internal modes dominated by recurrent prediction and memory-based processing, and external modes dominated by incoming sensory input^9,11,80^. This intrinsic rhythmic structure may provide the neural background against which spontaneous transitions become more or less likely to occur. In addition, against this backdrop of fluctuating neural states, changes in the relative strength of internal constraints and external input may further influence the likelihood of entering the transitional state. Spontaneous thought is considered to unfold under dynamic constraints from recent thoughts, affective state, goals, and current concerns^81,82^. When these constraints weaken, the ongoing internal trajectory may become less stable^81,83^, making cognition less committed to its current internal context and more susceptible to alternative inputs. Therefore, spontaneous transitions may be more likely to arise when intrinsic oscillatory cycles coincide with a shift in the balance between internal and external processing. Future work will be needed to directly test this account.

Although the think-aloud fMRI paradigm is well suited for tracking the neural correlates of spontaneous transitions between internal and external cognition, several methodological limitations warrant consideration. First, externally oriented thoughts comprised a small proportion of all reported spontaneous thoughts, yielding relatively few transitions involving external thoughts. This may have reduced the statistical power of the reported analyses^84,85^. Future work could address this issue by using longer scanning runs or multi-session designs to sample more thought transitions. Second, we were unable to examine the pre-transition neural responses uniquely associated with external-to-internal transitions. Because external thoughts were infrequent and brief, external-to-internal transitions often followed internal-to-external transitions in close temporal succession, causing their pre-transition hemodynamic responses to overlap substantially with the carryover signal from the preceding internal-to-external transition (Supplementary Fig. 1). Given that the two transition directions may be asymmetric^86^ and that characterizing both is therefore important for a comprehensive understanding of cognitive switching^3,14^, future studies should examine both directions using methods with higher temporal resolution than fMRI, such as electroencephalogram, magnetoencephalogram, or intracranial recordings. Third, the transition time identified from verbal responses may not precisely correspond to the actual moment of the underlying mental transition. Previous research suggests a temporal delay between the neural responses associated with thought generation and their subsequent verbalization^31,87^, and this delay may also vary across participants and individual transition events. Because the precise onset of spontaneous thought transitions remains inherently difficult to determine, interpretations of their temporal dynamics should be made with appropriate caution, and future studies should seek methods that can more accurately estimate transition timing. Finally, the external environment during scanning was relatively impoverished and stable. This may limit the generalizability of our findings to real-world settings, where rich and continuously changing external stimuli compete for attention. Future studies could employ more complex and dynamic sensory environments, for example using virtual reality paradigms^88,89^, to investigate internal-external thought transitions in more naturalistic contexts.

In conclusion, our study provides a multilevel account of the neural dynamics preceding naturally occurring transitions from internal representations to the external world, bridging research on spontaneous thought with work on cognitive reorientation. Specifically, the salience/ventral attention network, particularly the rTPJ, showed increased activation before internal-to-external transitions. This finding extends research on cognitive transitions beyond predominantly externally cued paradigms^2,3,20^, demonstrating that similar neural systems are recruited during spontaneous shifts. Exploratory brain-state analyses further suggest that this activation is preceded by a shift toward a flexible, transition-hub state^44^, extending the account to the level of large-scale brain dynamics and identifying such dynamics as a candidate system-level condition supporting spontaneous transitions. At the same time, the spatial correspondence between pre-transition activation and cholinergic receptor density provides a chemoarchitecturally grounded marker of cognitive reorientation and complements emerging neurochemical accounts of transitions between internal and external states^9,14^. More broadly, understanding how the brain flexibly regulates transitions between internal and external modes may also provide insight into everyday cognition and into clinical conditions characterized by excessive internal focus or distractibility, such as depression and attention-deficit/hyperactivity disorder^90–92^.

## METHODS

This study is a reanalysis of an existing think-aloud fMRI dataset^26^. Detailed descriptions of the dataset acquisition and annotation procedures are provided in previous publications^25,26^. All procedures were approved by the Institutional Review Board (IRB) of Johns Hopkins Medicine.

### Participants

The think-aloud dataset included a total of 118 healthy adults recruited from the Johns Hopkins University community. All participants were right-handed and fluent in English (aged 18-39 years), and reported normal hearing and normal or corrected-to-normal vision. Written informed consent was obtained from all participants. Following the criteria described in our previous study^25^, 43 participants were excluded due to poor-quality speech recordings, scanning interruptions, failure to follow instructions, excessive head motion, anatomical abnormalities, or MRI artifacts. An additional 19 participants were excluded from fMRI analysis in the current study due to the absence of internal-to-external thought transitions. The final sample included 56 participants (30 females, age 18-36 years, mean age 23.88 years).

### Procedures

Participants completed a 10-minute think-aloud task during fMRI scanning. They were instructed to continuously verbalize their spontaneous stream of thoughts, including but not limited to past memories, future plans, and ongoing perceptual experiences. Participants began speaking when the word “Begin” appeared on a gray screen. After 2 seconds, the cue was replaced by a white fixation cross, which remained on the screen for the remainder of the session. No other visual or auditory stimulus was presented. An MR-compatible microphone (FOMRI II; Optoacoustics Ltd.) was used to record participants’ speech. Detailed descriptions of the task instructions and experimental procedures are provided in our original study^25^.

### Behavioral data preprocessing

Audio recordings of participants’ think-aloud responses were transcribed either manually or automatically using Whisper (Large-v2 model; OpenAI) and subsequently corrected by human annotators. Each transcript was segmented into individual sentences, and the onset and offset times of each sentence were identified. Sentences consisting solely of filler utterances (e.g., “um”, “what else?”) were excluded.

For the present reanalysis, each remaining sentence was classified as reflecting either internal or external thought using the thought annotations included in the dataset, which were generated according to the procedures described in our previous publications^25,26^. In the original annotation, human annotators manually classified each sentence into one of several thought categories, including current experiences, episodic memory, future thinking, and semantic memory about oneself or the world and others. In addition, each sentence was rated on a 4-point Likert scale for the extent to which it expressed each sensory modality: vision, audition, olfaction, gustation, somatosensation, and interoception (1 = not at all, 4 = very much). These sensory modality ratings were generated using a large language model (Generative Pre-trained Transformer 5) and subsequently validated against human annotations for a subset of the transcripts.

We classified a sentence as externally oriented if it was labeled as a current experience and contained a report of sensory information (i.e., a sensory modality rating greater than 1) in at least one of the six modalities, including interoception. Although interoception refers to signals originating within the body, it was treated in the same manner as the other sensory modalities because, in our dataset, it primarily reflected low-level bodily experiences associated with the scanning context (e.g., "Um feels like my body is spinning"; "Now I’m actually getting kind of cold"). All remaining sentences were classified as internally oriented. These included a small proportion of sentences (M = 11.77%, SD = 11.92) labeled as current experiences but containing no reported sensory information (i.e., all six sensory modality ratings were equal to 1). These sentences primarily described thoughts about the think-aloud task itself (e.g., "I’m thinking about what the experiment is.") rather than externally derived sensory experiences.

Finally, consecutive sentence labels were used to classify thought transitions. For example, an internal-to-internal transition was defined as two consecutive internally oriented sentences, whereas an internal-to-external transition was defined as an internally oriented sentence followed by an externally oriented sentence.

### MRI data acquisition and preprocessing

MRI data were collected at Kennedy Krieger Institute on a Philips Ingenia Elition scanner (3T) with a 32-channel head coil. Functional images were acquired using a T2*-weighted multiband accelerated echo-planar imaging sequence (TR = 1.5 s, TE = 30 ms, flip angle = 52°, acceleration factor = 4, 60 oblique axial slices, voxel size = 2 × 2 × 2 mm^3^). Anatomical images were acquired with a T1-weighted MPRAGE pulse sequence (150 axial slices, voxel size = 1 × 1 × 1 mm^3^).

Preprocessing of anatomical images was carried out using FreeSurfer^93^. Functional image preprocessing was performed using fMRIprep^94^ with default settings, including correction for head motion and B0 magnetic-field inhomogeneity, coregistration to each participant’s anatomical image, and resampling to the MNI 152 volume space. Preprocessed functional images were spatially smoothed (FWHM = 4 mm). Nuisance regression was subsequently performed to regress out the six head-motion parameters, the mean cerebrospinal fluid and white matter signals, and second-order polynomial trends. The resulting time series were z-scored along the temporal dimension. For quality control, the first 10 volumes of the functional run were discarded. Motion outlier volumes (framewise displacement ≥ 1mm) were also excluded, together with one volume immediately preceding and following each outlier.

### Pre-transition activation analysis

For this and all subsequent fMRI analyses, we used the Schaefer 100-parcel cortical parcellation atlas^46^ to define cortical areas and functional networks as regions of interest (ROIs). The atlas divides the cerebral cortex into 100 spatially contiguous parcels (50 per hemisphere) based on resting-state functional connectivity. Each parcel was assigned to one of 17 large-scale functional networks identified in a previous study^95^. For parcel-wise analyses, preprocessed fMRI time series were averaged across voxels within each parcel for each participant. For network-wise analyses, parcel-wise activation time series were averaged across all parcels within each network, yielding activation time series for the 17 functional networks for each participant.

To examine neural activity preceding spontaneous thought transitions, we extracted parcel- and network-wise activation time series aligned to the onset of each post-transition thought for each participant. We analyzed two temporal windows around each transition while preserving the original transition onsets without applying temporal shifts to account for the hemodynamic response delay. First, our primary pre-transition analysis focused on a 3-TR (4.5 s) pre-transition window consisting of the two TRs immediately preceding the transition onset and the onset TR itself. This 4.5-s window is broadly consistent with prior work showing that spontaneous thought and awareness transitions are preceded by neural changes over several seconds (approximately 1-5 s)^31,47,48^. Two additional participants lacked valid data within this 3-TR pre-transition window because their corresponding TRs had been excluded during preprocessing, and were therefore excluded from this and all subsequent analyses using that window. Second, to visualize the broader temporal profile of brain activation surrounding transitions, we extracted a longer 21-TR (31.5-s) activation time course centered on the post-transition thought onset, spanning from 10 TRs before to 10 TRs after the onset TR.

For each participant and ROI, transition-locked activation time series were averaged across all transition events separately for internal-to-external and internal-to-internal transitions. Pre-transition activation for each transition type was then computed by averaging activity across the three TRs within the pre-transition window. Finally, two-tailed paired-samples t tests were performed at the group level for each ROI to compare pre-transition activation between internal-to-external and internal-to-internal transitions. Statistical significance was corrected for multiple comparisons across cortical parcels or functional networks.

### Controlling for semantic similarity

To test whether the effects of transition type on pre-transition neural activation were driven by differences in semantic relatedness between consecutive thoughts, we performed a control analysis (Fig. 2). Semantic relatedness at each thought transition was qualified by converting each think-aloud sentence into a text embedding using a pretrained language model (all-mpnet-base-v2) implemented in the Sentence Transformers Python module (https://www.sbert.net), following the approach used in our prior study^25^. Semantic similarity was then computed as the cosine similarity between the embedding vectors of the pre- and post-transition sentences.

Because internal-to-external transitions showed larger semantic shifts than internal-to-internal transitions (Fig. 2b), we constructed a semantically matched control set of internal-to-internal transitions for each participant. Specifically, for each participant, internal-to-internal transitions were ranked in ascending order of semantic similarity, and transitions with the lowest semantic similarity were selected first to form cumulative subsets (Fig. 2a). For each subset size, ranging from one transition to the full set of internal-to-internal transitions, we calculated the mean semantic similarity of the selected transitions and its absolute difference from that participant’s internal-to-external target mean. Near-best candidate subsets were then identified as those whose absolute differences fell within ±0.1 standard deviation of the minimum difference, where the standard deviation was computed across the absolute differences of all candidate subsets for that participant. One subset was then randomly selected from these near-best candidates. As a result, the final matched mean semantic similarity of the selected subset could be slightly lower or higher than the internal-to-external target mean. This procedure was repeated for all participants.

We then repeated the pre-transition activation analysis using the semantically matched subset of internal-to-internal transitions in place of the full sets used in the original analysis. For each parcel or functional network, mean activation during the pre-transition window was computed separately for internal-to-external transitions and the semantically matched internal-to-internal transitions within each participant. The two transition types were then compared at the group level using paired-samples *t*-tests.

### MRI-based eye movement analysis

Eye movement dynamics during scanning were estimated using DeepMReye with the default pretrained model^54^, a deep-learning model that decodes gaze position directly from the MR signal of the eyeballs (Fig. 3a). The model takes automatically extracted eyeball voxels as input and, using a three-dimensional convolutional neural network, predicts two-dimensional (horizontal and vertical) gaze coordinates for each functional volume. For each participant, DeepMReye generated 10 gaze-position estimates per fMRI volume. These estimates were median-averaged to obtain a single horizontal and vertical gaze-position estimate for each TR. TR-wise gaze displacement was then quantified as vector length (i.e., the Euclidean distance between consecutive gaze positions).

To examine eye movements around thought transitions, we extracted gaze vector length aligned to the onset of each post-transition thought for each participant. Based on previous studies indicating that changes in eye movements around event boundaries or cognitive switching occur within a relatively short interval (approximately 2-10 s around the transition)^52,96,97^, we analyzed a 5-TR (7.5-s) time window centered on the transition (i.e., from 2 TRs before to 2 TRs after the transition TR). For each participant, transition-locked gaze vector length time courses were averaged across all transitions separately for each transition type (internal-to-external vs. internal-to-internal). To test whether eye movements differed between transition types within the selected window, we conducted a two-way repeated-measures ANOVA with transition type and relative TR as within-subject factors. Post hoc paired-samples *t*-tests were then performed at each TR to compare the two transition types directly. Statistical significance was corrected for multiple comparisons across TRs.

To examine whether eye movements at thought transitions depended on the sensory modality of the upcoming external thought, internal-to-external transitions were categorized by the sensory modality of the post-transition thought, based on sentence-wise sensory modality ratings. A modality was considered present when its rating exceeded 1 (not at all present). To ensure each thought was attributed to a single sensory category, only thoughts with a rating above 1 on exactly one modality were included in the analysis. Because gustatory and olfactory internal-to-external transitions were identified in only one participant and no participants, respectively, these two modalities were excluded from analysis, leaving vision, audition, somatosensation, and interoception. For each of these four modalities, gaze vector length time courses were averaged across all corresponding transitions within each participant. At the time point of interest (i.e., the time point at which vector length during internal-to-external transitions, averaged across all modalities, was significantly greater than during internal-to-internal transitions), paired-samples *t*-tests then compared internal-to-external transitions in each modality separately against internal-to-internal transitions. Statistical significance was corrected for multiple comparisons across the four modality-specific tests.

To validate the quality of the DeepMReye-based eye tracking, we conducted a whole-brain analysis to examine whether gaze vector length was associated with activation in visual cortices (Supplementary Fig. 2), replicating previously reported findings^55^. For each participant, vector length was averaged across TRs within each thought sentence. The resulting values were z-scored across thoughts within each participant. Likewise, for each participant and cortical parcel, thought-wise activation was computed by averaging the preprocessed fMRI signal across TRs within each thought, with the temporal window for each thought shifted forward by 4.5 s to account for the hemodynamic response delay. For each parcel, we then fit a linear mixed-effects model predicting thought-level neural activation from mean gaze vector length, with participant included as a random intercept.

### Comparing different sensory modalities

To determine whether pre-transition activation was driven by a specific sensory modality of external thoughts, we grouped internal-to-external transitions according to the sensory modality of the post-transition thought (visual, auditory, somatosensory, or interoceptive), using the same procedure described for the eye movement analysis. For each participant, mean pre-transition activation was then computed separately for each sensory modality in salience/ventral attention network B and the right TPJ parcel. Because no participant had valid data for all four modality conditions, a repeated-measures ANOVA was not appropriate. Instead, we performed pairwise paired-samples *t*-tests between all available pairs of sensory modality conditions. Statistical significance was corrected for multiple comparisons across the six pairwise modality comparisons.

### Neurotransmitter receptor density analysis

To investigate the potential involvement of neurotransmitter systems, particularly the cholinergic system, in pre-transition activation during internal-to-external thought transitions, we examined the spatial correspondence between whole-brain maps of the pre-transition activation and neurotransmitter receptor density (Fig. 4). Based on prior work implicating both nicotinic and muscarinic acetylcholine receptors in the encoding of new information^17^, we selected two cholinergic receptors for analysis: the alpha-4-beta-2 nicotinic acetylcholine receptor (α4β2) and the muscarinic M1 receptor (M1). As control receptors, we selected the major excitatory and inhibitory receptors in the brain^58,59^: the N-methyl-D-aspartate receptor (NMDA) and the gamma-aminobutyric acid type A receptor (GABAA).

For each selected receptor, volumetric positron emission tomography (PET) receptor density maps were obtained from a publicly available dataset^42^ comprising neurotransmitter receptor maps derived from multiple previous PET studies^35–41^. All PET data were acquired from healthy adult participants. For each receptor, PET density maps were averaged across participants to create a group-level density map. The group-level receptor density map was then coregistered to standard MNI152 volumetric space and resampled to 2 mm resolution to match the Schaefer 100-parcel atlas used for parcellation^46^. We then computed the mean receptor density for each cortical parcel in the atlas by averaging voxel-wise density values within the parcel. Receptor density values were z-scored across parcels for each receptor.

Next, we performed a series of linear mixed-effects model analyses to examine the relationship between receptor density and pre-transition activation. First, separate models were fit for internal-to-internal and internal-to-external transitions, with pre-transition activation as the dependent variable, each receptor density as a fixed effect, and participant as a random intercept (e.g., internal-to-external pre-transition activation ∼ α4β2 + (1|participant)). Second, to directly test whether receptor-activation associations differed between transition types (internal-to-external vs. internal-to-internal), we fit models including receptor density, transition type, and their interaction as fixed effects (e.g., pre-transition activation ∼ α4β2 + transition type + α4β2 × transition type + (1|participant)). Third, to test whether each receptor explained unique variance in pre-transition activation when accounting for other neurotransmitter systems, we fit models including multiple receptors as fixed effects, separately for each transition type (e.g., internal-to-external pre-transition activation ∼ α4β2 + NMDA + GABAA + (1|participant)). The two cholinergic receptors were entered into separate models because they index the same neurotransmitter system.

### Large-scale brain state change

To test whether internal-to-external transitions were preceded by changes in large-scale brain states, we examined the similarity between whole-brain pre-transition activation patterns and brain state templates derived from a previous study^44^ (Fig. 5). In that study, a hidden Markov model was applied to fMRI data acquired during a variety of conditions, including attention tasks, movie watching, and rest, to identify recurring brain states across diverse cognitive and attentional contexts. Four states were identified: state 1 was characterized by prominent activation of the DMN and was associated with internally directed cognition and stable, sustained attention; state 2 showed dominant engagement of the DAN and was associated with externally directed attention and goal-oriented cognitive control; state 3 was characterized by increased activity in the somatomotor network and is thought to reflect sensorimotor information processing; state 4 exhibited relatively low amplitude activity across large-scale networks without a single dominant system and was interpreted as a baseline or transient state that bridges more specialized functional configurations.

The four whole-brain state templates, originally defined in MNI 152 space, were resampled to an isotropic voxel size of 2 mm to match the resolution of our functional data. For each state, a parcel-level activation template was generated by averaging voxel-wise activation values within each cortical parcel of the Schaefer 100-parcel atlas^46^. Pearson correlations were then computed between each parcel-level brain state template and the TR-by-TR parcel-level whole-brain activation patterns preceding thought transitions. Specifically, we analyzed an 11-TR window spanning the 10 TRs preceding the transition and the transition-onset TR. For each brain state, the resulting correlation time courses were first computed for individual transition events and then averaged across transitions of the same type (internal-to-external or internal-to-internal) within each participant. Group-level differences were assessed using a repeated-measures ANOVA with transition type and time point as within-participant factors. Post-hoc paired-samples *t*-tests compared pattern correlations between transition types at each time point for each brain state. Statistical significance was corrected for multiple comparisons across time points.

### Statistical tests

For comparisons between two conditions, we used paired-samples *t*-tests. For comparisons involving multiple within-participant conditions or factors, we used repeated-measures ANOVA. For linear mixed-effects models, fixed effects were evaluated using Wald tests. All statistical tests were two-tailed, and we reported raw (uncorrected) *p* values. Correction for multiple comparisons was performed using the Benjamini-Hochberg false discovery rate procedure. Participants with missing data in any condition were excluded from the corresponding analyses.

## DATA AVAILABILITY

Neuroimaging data are publicly available through the OpenNeuro repository (accession number: ds006067; version 2.0.0). Behavioral and annotation data are also publicly available through the Open Science Framework (https://osf.io/a56rm).

## CODE AVAILABILITY

The analyses in this study were conducted using publicly available MRI data processing software and Python packages. MRI preprocessing was performed using fMRIPrep (v23.2.1), and FreeSurfer (v7.2.0). Semantic similarity between verbal reports was calculated using Sentence Transformers (v5.1.0). Statistical analyses were performed using SciPy (v1.15.3) and statsmodels (v0.14.4).

## AUTHOR CONTRIBUTIONS

M.Zhao and H.L. conceived and designed the research. M.Zhao, M.Zhang, H.S., P.R.L., and H.L. analyzed the data. M.Zhao and H.L. wrote the original manuscript. M.Zhao, M.Zhang, H.S., and H.L. reviewed and edited the manuscript.

