## Supplementary for "Neural signatures of spontaneous transitions between internal and external thought"

**SUPPLEMENTARY FIGURES**

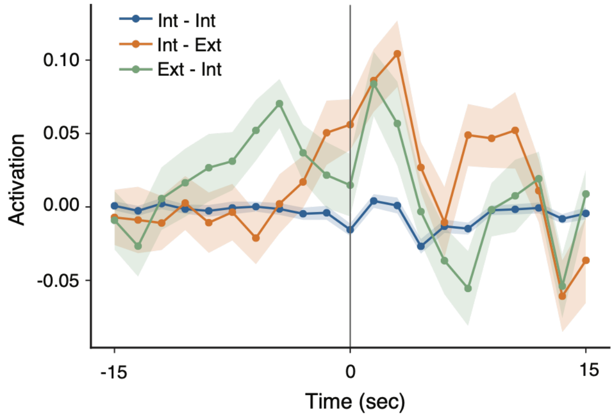


**Supplementary Fig. 1. Transition-locked salience/ventral attention network activation time courses for different types of thought transitions.** Mean activation time courses of salience/ventral attention network B are aligned to different types of thought transitions (blue = internal-to-internal, orange = internal-to-external, green = external-to-internal). Time zero represents the onset of the post-transition thought. Solid lines and circles indicate the mean activation across participants (N = 56) at each time point, and shaded areas indicate the SEM across participants. Notably, the peaks of the internal-to-external and external-to-internal time courses were separated by approximately 8 s, roughly matching the mean dwell time of external thoughts obtained from the 75 participants before participant exclusion (8.56 s). This timing suggests that the external-to-internal peak may partly reflect carryover activation from the preceding internal-to-external transition rather than a fully independent response to the return to internal thought.

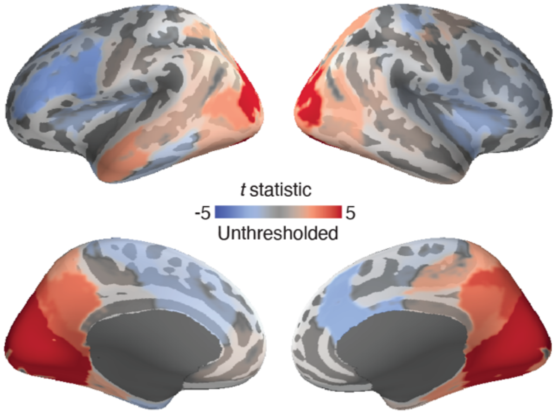


**Supplementary Fig. 2. Whole-brain activation associated with MRI-derived eye movement vector length.** Whole-brain *t*-statistic map of the relationship between brain activation and eye-movement vector length estimated from the fMRI data (N = 56). The maps are displayed on the lateral (top) and medial (bottom) surfaces of the left and right hemispheres of the inflated FreeSurfer template brain. Warm colors indicate positive associations between activation and vector length, whereas cool colors indicate negative associations. The color scale represents unthresholded *t* statistics.

### SUPPLEMENTARY TABLES

**Supplementary Table 1.** Internal-to-external versus internal-to-internal pre-transition activation contrast across the 17 functional networks.

| Direction of effect | Network | 95% CI | Cohen's *d* | *t* * | *p* (unc.) |
|---|---|---|---|---|---|
| Internal-to-external > Internal-to-internal | ControlA | [0.02, 0.11] | 0.37 | 2.67 | 0.009 |
| | ControlB | [-0.03, 0.05] | 0.09 | 0.64 | 0.526 |
| | ControlC | [-0.01, 0.08] | 0.21 | 1.53 | 0.133 |
| | DefaultB | [-0.01, 0.04] | 0.19 | 1.39 | 0.170 |
| | DorsAttnA | [-0.03, 0.07] | 0.13 | 0.93 | 0.358 |
| | DorsAttnB | [-0.02, 0.05] | 0.13 | 0.97 | 0.336 |
| | LimbicA | [-0.003, 0.05] | 0.24 | 1.77 | 0.082 |
| | SalVentAttnA | [0.01, 0.08] | 0.38 | 2.76 | 0.008 |
| | SalVentAttnB | [0.02, 0.08] | 0.44 | 3.27 | 0.002 |
| | SomMotA | [-0.01, 0.06] | 0.18 | 1.33 | 0.188 |
| | SomMotB | [-0.02, 0.07] | 0.15 | 1.12 | 0.267 |
| | TempPar | [0.001, 0.08] | 0.28 | 2.07 | 0.044 |
| | VisualA | [-0.04, 0.07] | 0.08 | 0.56 | 0.575 |
| | VisualB | [-0.05, 0.06] | 0.02 | 0.16 | 0.872 |
| Internal-to-internal > Internal-to-external | DefaultA | [-0.04, 0.03] | -0.07 | -0.54 | 0.591 |
| | DefaultC | [-0.08, -0.007] | -0.33 | -2.41 | 0.019 |
| | LimbicB | [-0.04, 0.01] | -0.14 | -1.02 | 0.313 |

*Degree of freedom = 53

**Supplementary Table 2.** Suprathreshold cortical parcels ($p < 0.05$, uncorrected) identified in the internal-to-external versus internal-to-internal pre-transition activation contrast.

| Direction of effect | Hemisphere | Network | Parcel | 95% CI | Cohen's *d* | *t* * | *p* (unc.) |
|---|---|---|---|---|---|---|---|
| Internal-to-external > Internal-to-internal | Left | ControlA | PFCl_1 | [0.03, 0.12] | 0.43 | 3.13 | 0.003 |
| | | | PFCl_2 | [0.01, 0.13] | 0.30 | 2.23 | 0.030 |
| | | SalVentAttnA | Ins_1 | [0.01, 0.12] | 0.30 | 2.20 | 0.032 |
| | | | Ins_2 | [0.02, 0.13] | 0.39 | 2.86 | 0.006 |
| | | | ParMed_1 | [0.02, 0.10] | 0.39 | 2.85 | 0.006 |
| | | | ParOper_1 | [0.004, 0.08] | 0.3 | 2.20 | 0.032 |
| | | SalVentAttnB | PFCl_1 | [0.01, 0.08] | 0.39 | 2.84 | 0.006 |
| | Right | ControlA | PFCl_1 | [0.01, 0.12] | 0.34 | 2.53 | 0.015 |
| | | | PFCl_2 | [0.02, 0.10] | 0.39 | 2.86 | 0.006 |
| | | DefaultB | PFCv_2 | [0.02, 0.12] | 0.39 | 2.87 | 0.006 |
| | | SalVentAttnA | Ins_1 | [0.03, 0.12] | 0.44 | 3.21 | 0.002 |
| | | SalVentAttnB | IPL_1 | [0.04, 0.15] | 0.51 | 3.76 | <0.001 |
| | | | PFCmp_1 | [0.01, 0.07] | 0.41 | 2.98 | 0.004 |
| | | TempPar | TempPar_2 | [0.003, 0.11] | 0.29 | 2.10 | 0.040 |
| | | | TempPar_3 | [0.001, 0.10] | 0.28 | 2.04 | 0.046 |
| Internal-to-internal > Internal-to-external | Left | DefaultC | Rsp_1 | [-0.15, -0.02] | -0.36 | -2.66 | 0.010 |
| | Right | DefaultC | Rsp_1 | [-0.12, -0.01] | -0.32 | -2.32 | 0.024 |

*Degree of freedom = 53

**Supplementary Table 3.** Internal-to-external versus internal-to-internal pre-transition activation contrast across the 17 functional networks after matching transitions for semantic similarity.

| Direction of effect | Network | 95% CI | Cohen's *d* | *t* * | *p* (unc.) |
|---|---|---|---|---|---|
| Internal-to-external > Internal-to-internal | ControlA | [0.02, 0.12] | 0.36 | 2.64 | 0.010 |
| | ControlB | [-0.03, 0.05] | 0.07 | 0.54 | 0.589 |
| | ControlC | [-0.003, 0.10] | 0.26 | 1.91 | 0.062 |
| | DefaultB | [-0.01, 0.04] | 0.15 | 1.13 | 0.265 |
| | DorsAttnA | [-0.03, 0.08] | 0.13 | 0.97 | 0.338 |
| | DorsAttnB | [-0.02, 0.06] | 0.16 | 1.16 | 0.252 |
| | LimbicA | [-0.01, 0.05] | 0.21 | 1.55 | 0.128 |
| | SalVentAttnA | [0.01, 0.09] | 0.39 | 2.85 | 0.006 |
| | SalVentAttnB | [0.02, 0.08] | 0.44 | 3.22 | 0.002 |
| | SomMotA | [-0.01, 0.07] | 0.23 | 1.69 | 0.098 |
| | SomMotB | [-0.01, 0.07] | 0.20 | 1.47 | 0.148 |
| | TempPar | [-0.003, 0.08] | 0.25 | 1.87 | 0.068 |
| | VisualA | [-0.04, 0.08] | 0.10 | 0.76 | 0.451 |
| | VisualB | [-0.04, 0.09] | 0.09 | 0.66 | 0.513 |
| Internal-to-internal > Internal-to-external | DefaultA | [-0.04, 0.03] | -0.03 | -0.25 | 0.802 |
| | DefaultC | [-0.07, 0.003] | -0.25 | -1.87 | 0.067 |
| | LimbicB | [-0.03, 0.01] | -0.16 | -1.18 | 0.242 |

*Degree of freedom = 53

**Supplementary Table 4.** Suprathreshold ($p$ < 0.05, uncorrected) cortical parcels identified in the internal-to-external versus internal-to-internal pre-transition activation contrast after matching transitions for semantic similarity.

<table>
<tr><th>Direction of effect</th><th>Hemisphere</th><th>Network</th><th>Parcel</th><th>95% CI</th><th>Cohen's d</th><th>t *</th><th>p (unc.)</th></tr>
<tr><td rowspan="13">Internal-to-external > Internal-to-internal</td><td rowspan="7">Left</td><td rowspan="2">ControlA</td><td>PFCl_1</td><td>[0.02, 0.12]</td><td>0.40</td><td>2.95</td><td>0.005</td></tr>
<tr><td>PFCl_2</td><td>[0.01, 0.13]</td><td>0.30</td><td>2.19</td><td>0.033</td></tr>
<tr><td rowspan="4">SalVentAttnA</td><td>Ins_1</td><td>[0.01, 0.12]</td><td>0.30</td><td>2.20</td><td>0.032</td></tr>
<tr><td>Ins_2</td><td>[0.03, 0.13]</td><td>0.43</td><td>3.13</td><td>0.003</td></tr>
<tr><td>ParMed_1</td><td>[0.02, 0.12]</td><td>0.39</td><td>2.88</td><td>0.006</td></tr>
<tr><td>ParOper_1</td><td>[0.01, 0.09]</td><td>0.33</td><td>2.48</td><td>0.016</td></tr>
<tr><td>SalVentAttnB</td><td>PFCl_1</td><td>[0.02, 0.09]</td><td>0.39</td><td>2.86</td><td>0.006</td></tr>
<tr><td rowspan="6">Right</td><td rowspan="2">ControlA</td><td>PFCl_1</td><td>[0.02, 0.14]</td><td>0.34</td><td>2.55</td><td>0.014</td></tr>
<tr><td>PFCl_2</td><td>[0.01, 0.11]</td><td>0.33</td><td>2.42</td><td>0.019</td></tr>
<tr><td>DefaultB</td><td>PFCv_2</td><td>[0.01, 0.12]</td><td>0.35</td><td>2.55</td><td>0.014</td></tr>
<tr><td>SalVentAttnA</td><td>Ins_1</td><td>[0.02, 0.12]</td><td>0.41</td><td>3.04</td><td>0.004</td></tr>
<tr><td rowspan="2">SalVentAttnB</td><td>IPL_1</td><td>[0.05, 0.16]</td><td>0.51</td><td>3.75</td><td><0.001</td></tr>
<tr><td>PFCmp_1</td><td>[0.01, 0.07]</td><td>0.38</td><td>2.80</td><td>0.007</td></tr>
<tr><td>Internal-to-internal > Internal-to-external</td><td>Left</td><td>DefaultC</td><td>Rsp_1</td><td>[-0.13, -0.01]</td><td>-0.30</td><td>-2.22</td><td>0.031</td></tr>
</table>

*Degree of freedom = 53

**Supplementary Table 5.** Pairwise comparisons of pre-transition activation in salience/ventral attention network B across sensory modalities.

| Pair | Mean difference | *df* | 95% CI | Cohen's *d* | *t* | *p* (unc.) |
|---|---|---|---|---|---|---|
| Auditory-Visual | -0.001 | 14 | [-0.14, 0.14] | -0.01 | -0.02 | 0.982 |
| Auditory-Somatosensory | -0.02 | 11 | [-0.18, 0.13] | -0.10 | -0.34 | 0.739 |
| Auditory-Interoceptive | 0.06 | 10 | [-0.12, 0.24] | 0.22 | 0.74 | 0.475 |
| Visual-Somatosensory | -0.05 | 13 | [-0.19, 0.09] | -0.20 | -0.20 | 0.471 |
| Visual-Interoceptive | 0.06 | 8 | [-0.06, 0.19] | 0.40 | 0.40 | 0.264 |
| Somatosensory-Interoceptive | 0.23 | 3 | [-0.14, 0.60] | 0.98 | 0.98 | 0.145 |

**Supplementary Table 6.** Pairwise comparisons of pre-transition activation in the right temporoparietal junction parcel across sensory modalities.

| Pair | Mean difference | *df* | 95% CI | Cohen's *d* | *t* | *p* (unc.) |
|---|---|---|---|---|---|---|
| Auditory-Visual | -0.12 | 14 | [-0.28, 0.04] | -0.41 | -1.60 | 0.132 |
| Auditory-Somatosensory | -0.05 | 11 | [-0.31, 0.21] | -0.11 | -0.40 | 0.698 |
| Auditory-Interoceptive | -0.003 | 10 | [-0.45, 0.44] | -0.004 | -0.02 | 0.988 |
| Visual-Somatosensory | -0.01 | 13 | [-0.2, 0.18] | -0.03 | -0.10 | 0.919 |
| Visual-Interoceptive | 0.07 | 8 | [-0.31, 0.46] | 0.15 | 0.44 | 0.670 |
| Somatosensory-Interoceptive | 0.10 | 3 | [-1.02, 1.22] | 0.14 | 0.28 | 0.798 |

**Supplementary Table 7.** Fixed-effect estimates from linear mixed-effects models testing receptor density × transition interactions on pre-transition activation.

| Receptor | Fixed effect | $\beta$ | 95% CI | $Z_{Wald}$ | $p$ |
|---|---|---|---|---|---|
| $\alpha_4\beta_2$ | Receptor density | 0.002 | [-0.002, 0.005] | 1.01 | 0.314 |
| | Receptor density × transition | 0.005 | [0.001, 0.010] | 2.23 | 0.026 |
| $M_1$ | Receptor density | -0.001 | [-0.004, 0.003] | -0.43 | 0.668 |
| | Receptor density × transition | 0.006 | [0.002, 0.011] | 2.63 | 0.008 |
| NMDA | Receptor density | 0.003 | [-0.001, 0.006] | 1.66 | 0.097 |
| | Receptor density × transition | -0.003 | [-0.008, 0.002] | -1.31 | 0.189 |
| $GABA_A$ | Receptor density | -0.0001 | [-0.003, 0.003] | -0.098 | 0.922 |
| | Receptor density × transition | -0.007 | [-0.012, -0.002] | -2.87 | 0.004 |

Note: The main effect of transition was identical across receptors: $\beta$ = 0.021, 95% CI = [0.017, 0.026], $Z_{Wald}$ = 8.93, $p$ <0.001.

**Supplementary Table 8.** Fixed-effect estimates from multiple-receptor linear mixed-effects models predicting pre-transition activation.

| Model | Predictor | $\beta$ | 95% CI | $Z_{Wald}$ | $p$ |
|---|---|---|---|---|---|
| Activation ~ $\alpha_4\beta_2$ + NMDA + $GABA_A$ + (1\|participant) | $\alpha_4\beta_2$ | 0.006 | [0.001, 0.011] | 2.37 | 0.018 |
| | NMDA | -0.0002 | [-0.005, 0.005] | -0.09 | 0.927 |
| | $GABA_A$ | -0.006 | [-0.011, -0.001] | -2.28 | 0.022 |
| Activation ~ $M_1$ + NMDA + $GABA_A$ + (1\|participant) | $M_1$ | 0.006 | [0.001, 0.010] | 2.56 | 0.010 |
| | NMDA | 0.002 | [-0.003, 0.006] | 0.68 | 0.498 |
| | $GABA_A$ | -0.008 | [-0.013, -0.003] | -3.28 | 0.001 |
